# Diet-induced metabolic and immune impairments are sex-specifically modulated by soluble TNF signaling in the 5xFAD mouse model of Alzheimer’s disease

**DOI:** 10.1101/2024.02.28.582516

**Authors:** Maria Elizabeth De Sousa Rodrigues, MacKenzie L. Bolen, Lisa Blackmer-Raynolds, Noah Schwartz, Jianjun Chang, Malú Gámez Tansey, Timothy Robert Sampson

## Abstract

Emerging evidence indicates that high-fat, high carbohydrate diet (HFHC) impacts central pathological features of Alzheimer’s disease (AD) across both human incidences and animal models. However, the mechanisms underlying this association are poorly understood. Here, we identify compartment-specific metabolic and inflammatory dysregulations that are induced by HFHC diet in the 5xFAD mouse model of AD pathology. We observe that both male and female 5xFAD mice display exacerbated adiposity, cholesterolemia, and dysregulated insulin signaling. Independent of biological sex, HFHC diet also resulted in altered inflammatory cytokine profiles across the gastrointestinal, circulating, and central nervous systems (CNS) compartments demonstrating region-specific impacts of metabolic inflammation. In male mice, we note that HFHC triggered increases in amyloid beta, an observation not seen in female mice. Interestingly, inhibiting the inflammatory cytokine, soluble tumor necrosis factor (TNF) with the brain-permeant soluble TNF inhibitor XPro1595 was able to restore aspects of HFHC-induced metabolic inflammation, but only in male mice. Targeted transcriptomics of CNS regions revealed that inhibition of soluble TNF was sufficient to alter expression of hippocampal and cortical genes associated with beneficial immune and metabolic responses. Collectively, these results suggest that HFHC diet impairs metabolic and inflammatory pathways in an AD-relevant genotype and that soluble TNF has sex-dependent roles in modulating these pathways across anatomical compartments. Modulation of energy homeostasis and inflammation may provide new therapeutic avenues for AD.

**Highlights:**

- HFHC diet triggers metabolic and inflammatory dysregulation in 5xFAD mice
- Aβ42 and Aβ40 increase in the CNS of male 5xFAD mice fed a HFHC diet.
- SolTNF inhibition partially reverts HFHC diet-induced metabolic and inflammatory dysregulation in male, but not female, 5xFAD mice.

## Introduction

Alzheimer’s disease (AD) is the most common age-related cause of dementia, with the late-onset type (LOAD) becoming increasingly prevalent (1, 2). Pathologically, AD is characterized by the accumulation of amyloid-β protein (Aβ) and hyperphosphorylated tau deposition within neurons of the central nervous system (DOI: 10.7150/ijbs.57078). Strong evidence suggests that genetic mechanisms alongside environmental and lifestyle factors contribute to pathological protein deposition in AD, ultimately associated with neuroinflammation, neuronal death, and cognitive impairment (3–5).

Immune dysregulation, particularly neuroinflammation, is a central observation in AD and various animal models. In both human incidences and experimental models, amyloid deposits promote an inflammatory cytokine and chemokine response in the brain (6–8). While this cascade may be initially beneficial to control proteinopathy, the prolonged activation of glial cells can aggravate chronic neuroinflammation--a vicious cycle that accelerates disease progression and can contribute to further protein accumulation and neurodegeneration (9, 10). Systemic inflammatory conditions, including metabolic syndrome, is one risk factor for AD (11). However, the molecular and signaling mechanisms that underlie this relationship are not yet fully understood (12, 13). Elevated circulating cytokines, such as tumor necrosis factor (TNF), are associated with both increased risk of AD and with insulin resistance (14–16). TNF is cleaved into a soluble form (solTNF), that preferentially mediates inflammatory processes and exacerbating the hallmarks of AD pathology, such as β-secretase 1 expression in the CNS, the production of toxic Aβ peptides (Aβ 40 and Aβ 42), activation of microglia, and the release of other inflammatory mediators (17–19). Systemically, increased TNF is associated with comorbidities of metabolic dysregulation, including insulin impairment, vascular dysfunction, and dyslipidemia that further increases risk of AD (20–22).

Obesity and dementia have established sex-dependent outcomes in disease risk and progression in both conditions (23–27). Recent studies suggest that the sex-specific disparities in AD may arise due to unique metabolic and immune responses between the sexes, highlighting the potential role of environmental risk factors on inflammation (28–31). Diet is one modifiable factor that strongly impacts metabolic and immune responses across body systems (32, 33). Diets rich in specific fatty acids and sugars can lead to increased intestinal permeability and circulation of intestinal microbe-derived molecules with various effects on host physiology (34, 35). Some of these bacterial components elicit inflammatory responses that can become chronic (36–38). Specifically, diets high in fat and carbohydrates (high-fat, high-carbohydrate; HFHC) increase the translocation of bacterial lipopolysaccharides, which strongly activate circulating lipopolysaccharide-binding protein (LBP) and trigger Toll-like receptor (TLR4)-mediated inflammation (39). Blood-brain-barrier (BBB) dysfunctions are also associated with an increased systemic inflammatory tone. BBB dysregulation as a result of chronic inflammation, can be a result of signaling by peripheral TNF, allowing circulating immune cells to infiltrate into the parenchyma (40). In line with this, prior work from our team demonstrated that an HFHC diet impacts both the immune cell profile in circulation and within the CNS of the 5xFAD mouse model of AD (37).

Here, we employed a diet-induced model of metabolic syndrome in the 5xFAD mouse model to determine the effects of solTNF inhibition on central and peripheral metabolic and inflammatory responses that impact AD. We identify sex-dependent outcomes of HFHC on inflammatory cytokine responses across immune compartments, including in circulation, and within intestinal and CNS tissues. We further note that inhibition of solTNF reversed specific metabolic and inflammatory outcomes in the CNS of male animals, but not female animals. Thus, our data indicate that while HFHC diet detrimentally impacts many parameters in 5xFAD mice, solTNF signaling has sex-specific effects to limit these pathologies.

## Materials and methods

### Animal husbandry

Female and male 5xFAD mice, on a congenic C57Bl/6J background (Jackson Labs: #034848) were maintained by crossing with C57Bl/6J wildtype mice. Mice were co-housed (2-5 per cage) with mixed genotype littermates in sterile, microisolator cages, containing plastic igloos for enrichment under a 12 h light/dark cycle with ad libitum access to food and drinking water. Mice were genotyped with the following primers, using parameters from the commercial vendor: APP Forward 5’-AGGACTGACCACTCGACCAG-3’, APP Reverse 5’-CGGGGGTCTAGTTCTGCAT-3’; PS1 Forward 5’-AATAGAGAACGGCAGGAGCA-3’, PS1 Reverse 5’-GCCATGAGGGCACTAATCAT-3’. All animal experiments were approved by the Institutional Animal Care and Use Committee of Emory University (PROTO201900056) and conducted in compliance with the National Institute of Health Guide for the Care and Use of Laboratory Animals.

### Diet-induced obesity

Eight-week old 5xFAD female and male mice were fed a control diet (CD) (#TD.7001, 4% kcal fat, ENVIGO plus regular sterile drinking water) or a HFHC diet (#TD.88137 42% kcal fat, ENVIGO, plus 30% fructose (#F012, Sigma-Aldrich) w/v in sterile drinking water) for 8 weeks (Diet details found in Support information). Mice were weighed weekly. Food was replaced twice weekly, and 30% sterile fructose drinking water was replaced daily. All mice used in the diet studies were randomly assigned to the following groups: Control diet (CD); high-fat high carbohydrate diet (HFHC). Mice were kept on their respective diets for the duration of the study.

### Targeting solTNF signaling

After 4 weeks of consumption of a CD or HFHC diet, female and male 5xFAD mice (n=10-14) received subcutaneous (*sq*) injections of either the brain-permeant biologic XPro1595® (10mg/kg) or vehicle (sterile saline-1ml/kg) twice a week for the remaining 4 weeks of diet intervention. The XPro1595 dose was selected based on previously published work (41) and kindly provided by Dr. David Szymkowski (Xencor, Inc). For diet and soluble TNF studies, mice were randomly distributed into the groups: Control diet saline (CDS); high-fat high-carbohydrate diet saline (HFHC S) and high-fat high-carbohydrate diet XPro1595 (HFHC XPro).

### Fecal output and water content

Animals were gently removed from their home cages and individually placed in clear sterile 1L plastic beakers. Each beaker was covered with a paper towel during the test. The fecal pellets found were counted in 5min intervals. The number of pellets present in the beaker floor/wall were cumulatively counted and recorded up to 30min. The mice were returned to their home cages and the pellets collected were used for fecal water content analysis. For water content assay, after recording empty 1.5mL tubes weight, fresh fecal pellets per mouse was collected in each tube and the weight of tube plus feces were recorded. The microcentrifuge tubes and fecal contents were immediately placed on heat block set to 90C-100°C, under a fume hood and incubated for 24hrs to remove water from the fecal samples. After 24hrs the tubes containing the dry pellets were weighed. The percent of water content was calculated as following: Full tube weight – Empty tube weight = wet weight of fecal pellets. Dry tube weight – Empty tube weight = dry weight of fecal pellets. Percent water content= (Wet weight – dry weight) / wet weight.

### Tissue collection

At 16 weeks of age mice were humanely euthanized under isoflurane anesthesia, blood was collected by cardiac puncture into 7.5% EDTA Coviden tubes and immediately centrifuged at 10k x*g* for 12min at 4°C. Plasma was collected and stored at -80°C. Following exsanguination, mice were cervically dislocated. The gonadal fat pad, liver, cecum and spleens were removed and weighed. Colon and small intestine were measured. The hippocampus, cortex and approximately 4mm sections of clean proximal colon were flash frozen in liquid nitrogen and stored at -80°C. A subset of mice was transcardially perfused with ∼40–60mL of saline (0.9% NaCl: 25°C, pH 7.4). Left brain hemispheres were placed in 15mL tube filled with 8mL of a 4% paraformaldehyde in PBS solution, overnight, for fixation at 4°C. The brains samples were then transferred to tubes contain 8mL of 30% sucrose solution for dehydration at 4°C. After dehydration, the brain hemispheres were embedded in optimal cutting temperature compound (OCT) prior to frozen sectioning and stored in -80°C.

### Molecular assessments

For multiplexed immunoassays, plasma was diluted 1:4 using the MSD homogenization buffer (1% Triton-X100, 1M Tris HCL, 0.5M MgCl_2_ and 0.1M EDTA-pH 7.8). Cortex, hippocampus and colon tissue samples were sonicated in 1X MSD buffer on ice and the protein concentration was determined by Pierce BCA Protein Assay Kit (#23225, Pierce Scientific). Protein lysates (∼60-65µL at 1.6µg/µl) were used for immunoassays that were carried out per manufacturer’s instructions on the (QuickPlex SQ 120MM, Meso Scale Diagnostics) and analyzed on the MSD Discovery workbench 4.0 platform. Plasma insulin and leptin were measured using the Mouse Metabolic Kit (MSD # K15124C-1). The Mouse V-PLEX Proinflammatory Panel 1 Mouse Kit (#K15048D-1), Proinflammatory 7-Plex Ultra-Sensitive Assay (#K15012C-1), 19-Plex MSD kit (# K15255D-1), (U-PLEX Mouse IL-13 Assay (#K152UBK-1) were used to assess inflammation in the plasma, proximal colon, cortex and hippocampus. The V-PLEX Aβ Peptide Panel 1 (6E10) Kit (5 Plate) (MSD # K15200E-2) was used to assess soluble amyloid concentrations in the cortex, hippocampus and colon.

Measurements for plasma cholesterol (Kit #MAK043, Sigma-Aldrich, St. Louis, MO), glucose (#ab65333, Abcam) and LBP (#ab269542, Abcam), and proximal colon and plasma lipocalin 2 (LCN2) (#MLCN20, R&D Systems Minneapolis, MN) were also performed according with the manufacturer’s instructions. For colon LCN2 proximal colon tissue was homogenized by sonication with phosphate buffered saline (PBS) containing 0.1% Tween20 and centrifuged for 10min at 3k x*g* at 4°C. Supernatants were collected and total protein was diluted to a final concentration of 100mg/mL. Synergy 4 (BioteK) plate reader was used to determine the concentration of LCN2, LBP, glucose and cholesterol. Plasma and tissue assays were performed in duplicate by an experimenter blinded to the identity of samples. Western blot analyses were performed as previously described (22). Briefly, flash frozen cortex, hippocampus and proximal colon samples were homogenized by sonication in a 1X MSD buffer containing protease and phosphatase inhibitors (#A32961, Peirce, Thermo Fisher). After sonication, samples were centrifuged at 13k x*g* for 20min at 4°C. Protein concentration of the supernatant were determined by Pierce BCA Protein Assay Kit (Pierce Scientific #23225). Samples were diluted to 1µg/µl in 2X Lamelli buffer (BioRad #1610747) and boiled at 90°C for 5min. Samples were resolved using SDS-PAGE, transferred overnight to PVDF membranes at 4°C, and blocked with 5% BSA in 1X Tris-buffered saline (20 mM, Tris pH 7.5, 150 mM NaCl) and 0.1% Tween20) for 1 h. Membranes were probed overnight at 4°C with indicated primary antibodies at the concentrations listed in Supplementary Table S2. After incubation with the indicated horseradish peroxidase (HRP)-conjugated secondary antibody (1:1000), bands were visualized with SuperSignal™ West Femto Maximum Sensitivity Substrate (#34094, Thermo Fisher) and protein band optical intensity was measured using Azure Imaging System and densitometric analysis was performed using Image Studio Lite quantification software. The densities of the phosphorylated protein bands were measured relative to the targeted total protein levels. Densitometric values were normalized relative to β-actin levels from the same sample.

### Microbiome profiling

Fresh fecal pellets from female 5xFAD mice were collected and immediately frozen from baseline time point, after 4 weeks and 8 weeks of CD or HFHC diet intake and of 8 weeks of diet consumption and 4 weeks of XPro1595/saline treatment. Fecal samples were stored at −80^◦^C before bacterial DNA was extracted, quantified, purified and 16rRNA sequencing performed. The samples were processed and analyzed with the ZymoBIOMICS® Targeted Sequencing Service (Zymo Research, Irvine, CA). Additional details found at supplemental information file.

### RNA isolation and gene expression analysis

RNA was extracted from frozen cortex and hippocampus of male and female 5xFAD mice (CDS, HFHCS and HFHC XPro groups). The Qiagen RNeasy kit (# NC9307831, Qiagen) was followed per manufacturer’s instructions and Trizol RNA isolation was completed. RNA concentration and quality were identified via Qubit and Quant-it Assay (#Q33263) and Agilent 2100 Bioanalyzer (#G2939A) according to manufacturer’s instructions. The NanoString nCounter® Analysis System (NanoString Technologies, Seattle, USA) was used to complete the mouse neuropathology nCounter®assay (XT-CSO-MNROP1-12), according to manufacturer’s instructions. In brief, 100ng RNA in 5ul H20 from fresh-frozen tissue was loaded per sample. RNA was hybridized overnight at 65°C with biotin-labeled capture probes and fluorescently labeled reporter probes. Post hybridization, samples were pipetted into the nCounter chip via the nCounter MAX/FLEX robot where non-adhered capture and reporter probes were removed and hybridized mRNAs were immobilized for imaging. Raw count RCC files were normalized on the nSolver analysis software v4.0 (NanoString Technologies) according to manufacturer’s instructions and exported into ROSALIND® where built in statistical analysis was completed per manufacturer’s instruction. Normalization of all gene expression was preformed using NanoString nSolver v4.0 default settings, where raw counts were normalized to the geometric mean of positive/negative controls and housekeeping probes within the panel. Probes were not selected for analysis if greater than half of the samples do not meet sample specific threshold, as a result said pruned probes produce a log2 Fold Change of 0 and an adjusted p value of 1.

### Statistical Analysis

As indicated in the figure legends, data are expressed as mean ± s.e.m. Statistical significance was determined using two-tailed unpaired Student’s t test to compare two experimental groups and One-way analyses of variance (ANOVA) followed by Tukey’s or Sidak’s post hoc tests. A *p* value < 0.05 was considered to be significant. For Nanostring analysis, the fold change of 1.5 or greater was considered significant. Adjusted *p* values represent the Benjamini-Hochberg method of false discovery rate (FDR) corrected *p*-values. Statistical analysis was conducted using GraphPad Prism 8.

## Results

### High-fat, high carbohydrate diet promotes diet-induced insulin impairment in 5xFAD mice

Our team previously identified that HFHC diet resulted in AD-relevant metabolic impacts in wild-type mice (22). We therefore sought to establish whether HFHC diet could interact with an AD genetic model to exacerbate metabolic, immune, and amyloid pathologies. Recent work from our team indicated only mild insulin impairment in 5xFAD mice when provided high fructose solely in solid food (42). Therefore, 2-month old female 5xFAD mice were provided a high-fat, high-sucrose solid food diet and fructose-replete drinking water *ad lib* for 8 weeks. Following dietary intervention, we measured central morphometric parameters established for diet-induced obesity. Similar to prior work (43–45), HFHC-fed 5xFAD mice exhibited increased body weight gain, gonadal fat and liver weight when compared to CD group (Fig. 1A-C). Insulinemia, leptinemia and hypercholesterolemia was also apparent (Fig. 1D-F). Circulating lipocalin-2 (LCN2), a downstream TNF inflammatory molecule associated with insulin impairment was also significantly elevated in plasma HFHC-fed mice (Fig. 1G). Overall demonstrating that HFHC induces significant, systemic metabolic dysregulation in this mouse model of AD-related pathology.

**Figure 1.**
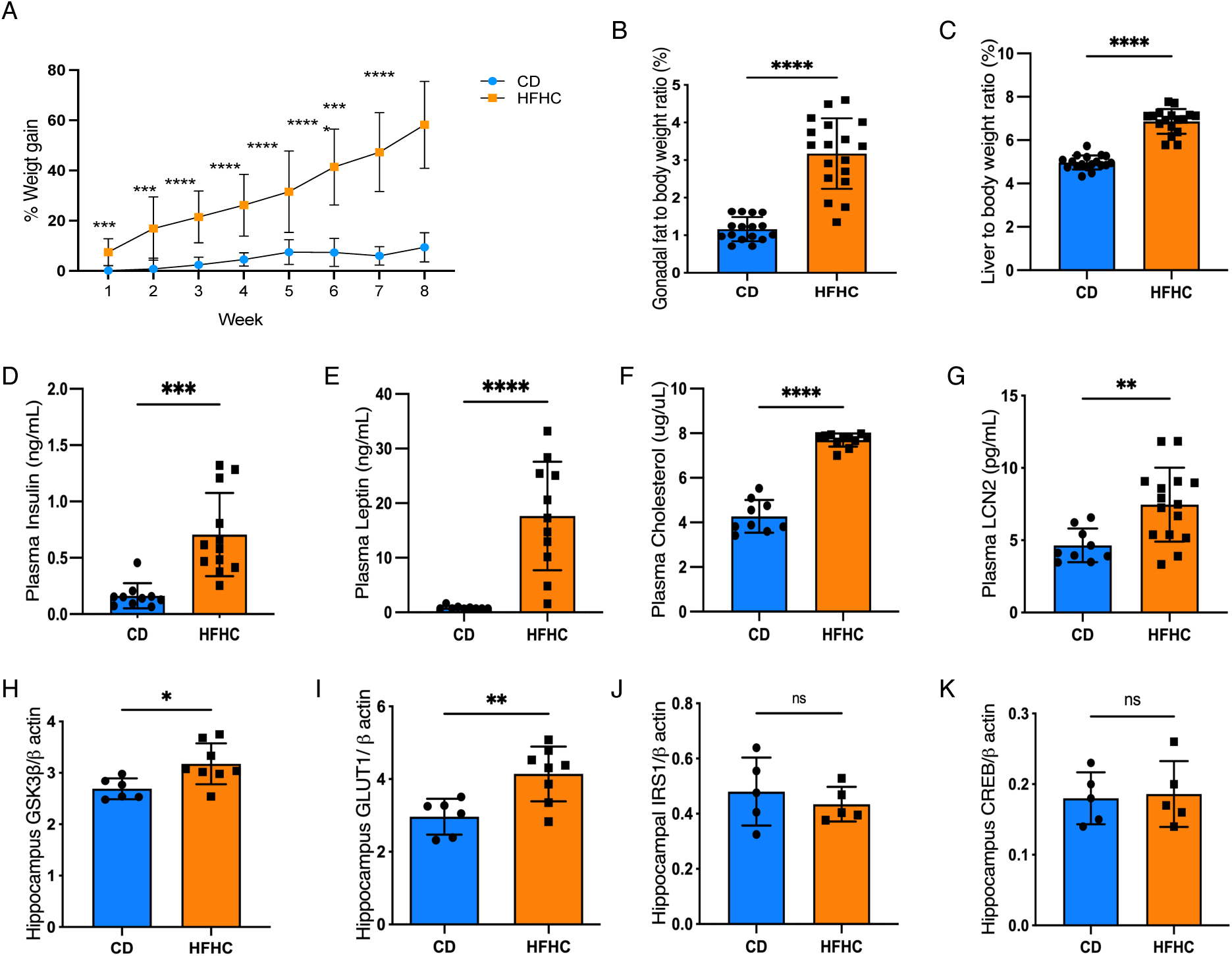
High-fat high-carbohydrate diet promotes systemic and hippocampal metabolic alterations in 5xFAD mice. (**A**) Weekly percentage of body weight gain following HFHC diet. (**B**) Gonadal adipose tissue and (**C**) liver weights after 8 weeks of a CD or HFHC intake respectively. (**D-G**) Circulating insulin, leptin, cholesterol and LCN2 of 5xFAD mice at 8wks by multi-plex ELISA. (**H-K**). Protein abundances of hippocampal GSK3b, GLUT1, IRS1 and CREB by western blot after diet treatment. Points represent individuals, bars represent the mean and SEM. Data assessed by two-way ANOVA with repeated measurements, followed by Sidak’s post hoc in (**A**) or unpaired, two-tailed Student’s t-test in (**B-K**). \**p* ≤0.05, \*\**p* ≤ 0.01, \*\*\**p* ≤ 0.001, \*\*\*\**p* ≤ 0.0001, ns=non-significant. Full western blots available in the supplementary data.

Given our observations of systemic insulinemia in the HFHC-fed 5xFAD mice, we next investigated whether diet-induced obesity in this animal model of AD would impact pathways which contribute to insulin/glucose impairment in the CNS. Glycogen synthase kinase-3 beta (GSK3β) and its downstream signaling factors, AKT and CREB have roles in insulin/glucose signaling, inflammation, and AD pathology (46–48). We observed an elevation of both GSK-3β and glucose transporter 1 (GLUT1) protein in the hippocampus of HFHC-fed 5xFAD mice, without any observable effect of diet on insulin receptor substrate 1 (IRS1) (Fig. 1H-K). Further in-depth examination of insulin-dependent pathways in this tissue surprisingly revealed that HFHC diet had no impact on production of insulin receptor alpha (IRα), insulin receptor beta (IRβ), GLUT3, and phospho-serine 473 AKT (pAKT Ser473) (Supplemental Fig. S1A-D). There were also no significant differences on hippocampal production of CREB or phosphorylation of GSK-3β (at sites serine 9 and 389, involved in its inactivation) following diet consumption (Supplemental Fig. S1E-H). Overall, this suggests that at this time point, HFHC consumption in 5xFAD mice promotes an adaptive response to the increase of energy supply, rather than severe insulin impairment in the brain. In the frontal cortex, we observed no effect of HFHC diet on GSK-3β, GLUT1, GLUT3, IRS1, and CREB (Supplementary Fig. S1I-L), demonstrating region-specific differences in HFHC impacts in the CNS. There similarly was no effect of HFHC diet on the pre-synaptic marker synaptophysin nor post-synaptic PSD95 (Supplemental Fig. S1M, N), indicating that HFHC did not accelerate synaptic dysregulation in the 5xFAD model. Further we do not observe any significant impacts on the abundance of Aβ38, 42, 40 or their ratios in either hippocampal or cortical tissue (Supplemental Fig. S2). Together, these findings indicate that HFHC-induced systemic metabolic inflammation in young 5xFAD mice is associated with GSK-3β overexpression, rather than insulin impairment or amyloid deposition in the CNS.

### Diet-induced insulin impairment promotes compartment-specific inflammation between the GI tract and brain of 5xFAD female mice

Diet-induced obesity is well-established to trigger chronic systemic inflammation. We noted that the HFHC induced a systemic inflammatory profile in 5xFAD mice, as measured by increases in circulating TNF, CCL2, CXCL10 and CXCL2, and decreases in plasma IL-2 and IL-33 compared to CD-fed 5xFAD animals (Fig. 2A-B). In the GI tract, we observed a pro-inflammatory profile as well, but characterized by different patterns of local cytokines. Colonic tissue of HFHC-fed 5xFAD mice had elevated IL-1ß, CXCL1 and CXCL2 and decreased IL-5 (Fig. 2C-D). Regional differences in cytokine profiles within brain tissue was also observed. In the cortex, we noted an increase in cortical CXCL2 and CCL3 and a decrease in IL-27p28, IL-5 and IL-10 in HFHC-fed 5xFAD mice. However, in hippocampal tissue of the same animals, only IL-1ß and IL-13 were increased (Fig. 2E-F). Despite the HFHC-induced increase of IL-1ß within hippocampus of 5xFAD mice, we did not observe elevated caspase 1, cleaved caspase-1, or NLRP3 (Supplemental Fig. S1H, O, P) in this specific brain region), suggesting that HFHC does not induce an overall cytotoxic environment in these brain regions of 5xFAD mice. Interestingly, the CNS cytokine profile observed in the female HFHC-fed 5xFAD female mice does not mirror the cytokine profile seen in circulation. Thus, HFHC differentially impacts immune signaling in each of these compartments, and is not simply derived from circulating cytokines crossing into the parenchyma.

**Figure 2.**
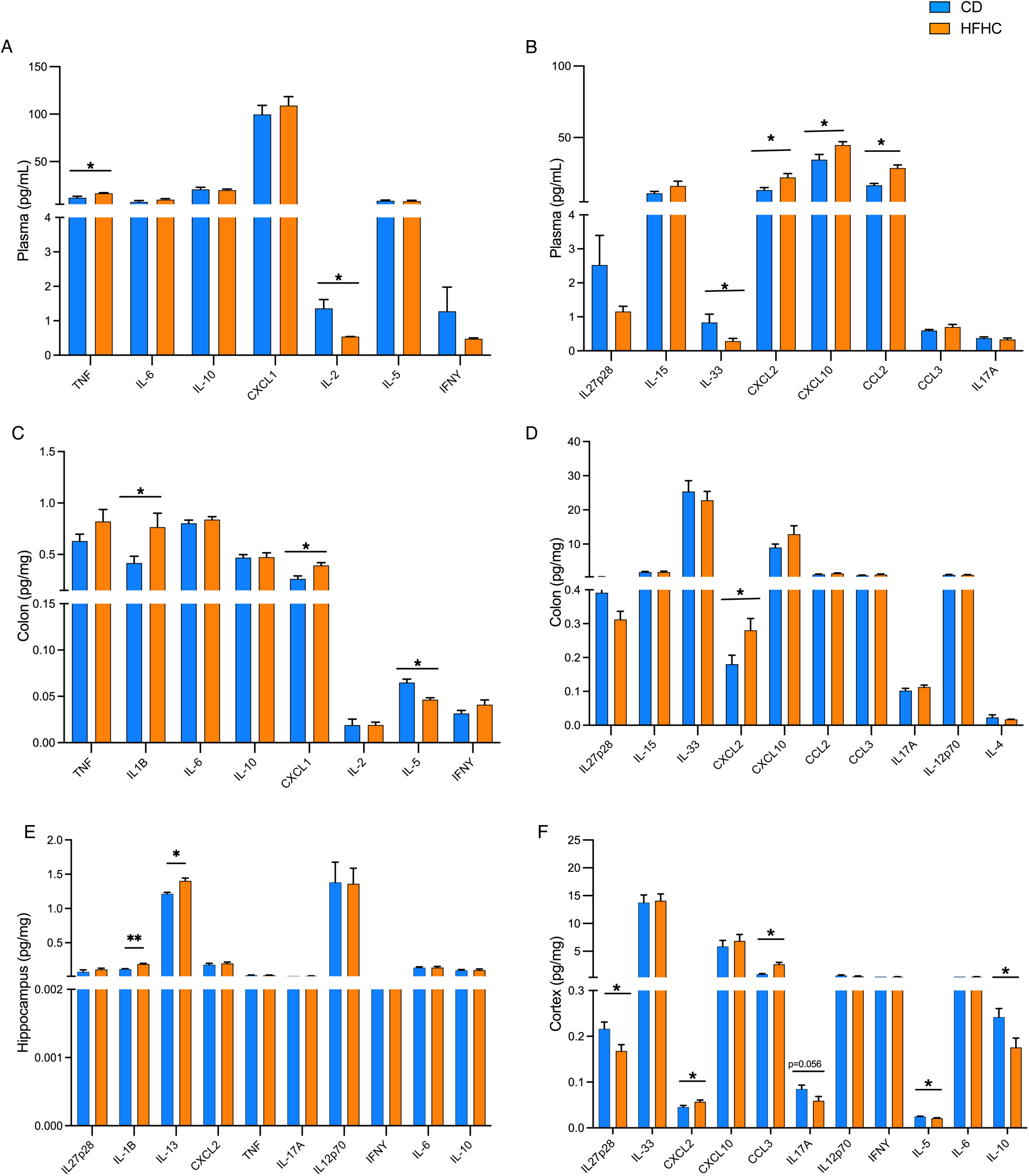
HFHC diet triggers compartment-specific immune responses across the brain and gut of 5xFAD mice. Cytokine and chemokine concentrations assessed by multiplexed ELISA in the (**A, B**) plasma, (**C, D**) colon, (**E**) hippocampus and (**F**) cortex of 5xFAD female mice fed 8 wks of CD or HFHC diet. Bars represent mean and SEM. Data assessed by two-tailed T test, adjusted with a false discovery rate (FDR) correction using a threshold of 1% for multiple comparisons (Benjamini, Krieger, and Yekutieli method). \**p* ≤ 0.05, n=5-12.

Signals derived from the GI tract itself are thought to contribute to diet-induced immune alterations. We found that HFHC diet decreased the length of the colon and small intestine and lowered cecal weight in 5xFAD mice (Supplemental Fig. S3A-C). The production of the tight junction protein occludin in the colon was decreased in the HFHC-fed mice, without significant alterations to levels of zona occludens 1 (ZO-1) and claudin-2 (Supplemental Fig. S3D-F). This marked decrease in colonic occludin suggests an increase in the permeability of the intestinal barrier, which may promote translocation of bacterial products and impact systemic immune responses across compartments. These microbial products are largely derived from the resident gut microbiome. Since dietary interventions such as HFHC diet, can shape the gut microbiome community, we assessed the community composition by 16S rRNA sequencing analysis. Both alpha and beta diversity metrics showed dissimilarity between fecal microbiome samples at 8 weeks of HFHC diet intake when compared to communities assessed at baseline and 4 weeks after the start of the dietary intervention (Supplemental Fig. S4A-D), whereas 5xFAD mice fed the CD displayed no community shift over time (Supplemental Fig. S4A). Within the 5xFAD mice, HFHC diet resulted in an enrichment of *Alcaligenaceae*, *Prevotellacea,* and *Bacteriodales* families and a decrease of *Christensenellaceae, Porphyromonadaceae, Streptococcaceae, Enterococcaceae, Clostridiaceae,* and *Desulfovibrionaceae* families (Supplemental Fig. 4E). Together, these data support the notion that HFHC promotes a highly altered intestinal environment in 5xFAD mice, through immune signaling, barrier permeability, and microbiome composition.

*Soluble TNF signaling does not mitigate HFHC-induced morphometric alterations in 5xFAD mice* TNF, elevated in circulation of HFHC-fed, 5xFAD mice, is an adipokine that regulates indirectly body energy metabolism. Given its role in metabolic inflammation, we next sought to determine whether solTNF signaling contributed to HFHC-induced obesity in 5xFAD mice. One month following HFHC or CD diet intervention, the brain-permeant solTNF inhibitor XPro1595 or vehicle (saline) were dosed twice weekly subcutaneously for 4 weeks in both female and male 5xFAD mice, independently. Diet-induced morphometric parameters, including weight gain, gonadal and liver weights were not affected by XPro1595 intervention (Fig. 3A-H). Colon and small intestine length remained shortened with no impact of solTNF inhibition (Fig. 3I-L), with little impact on physiological intestinal functions as measured by fecal output and fecal water content (Fig. 3M-P).

**Figure 3.**
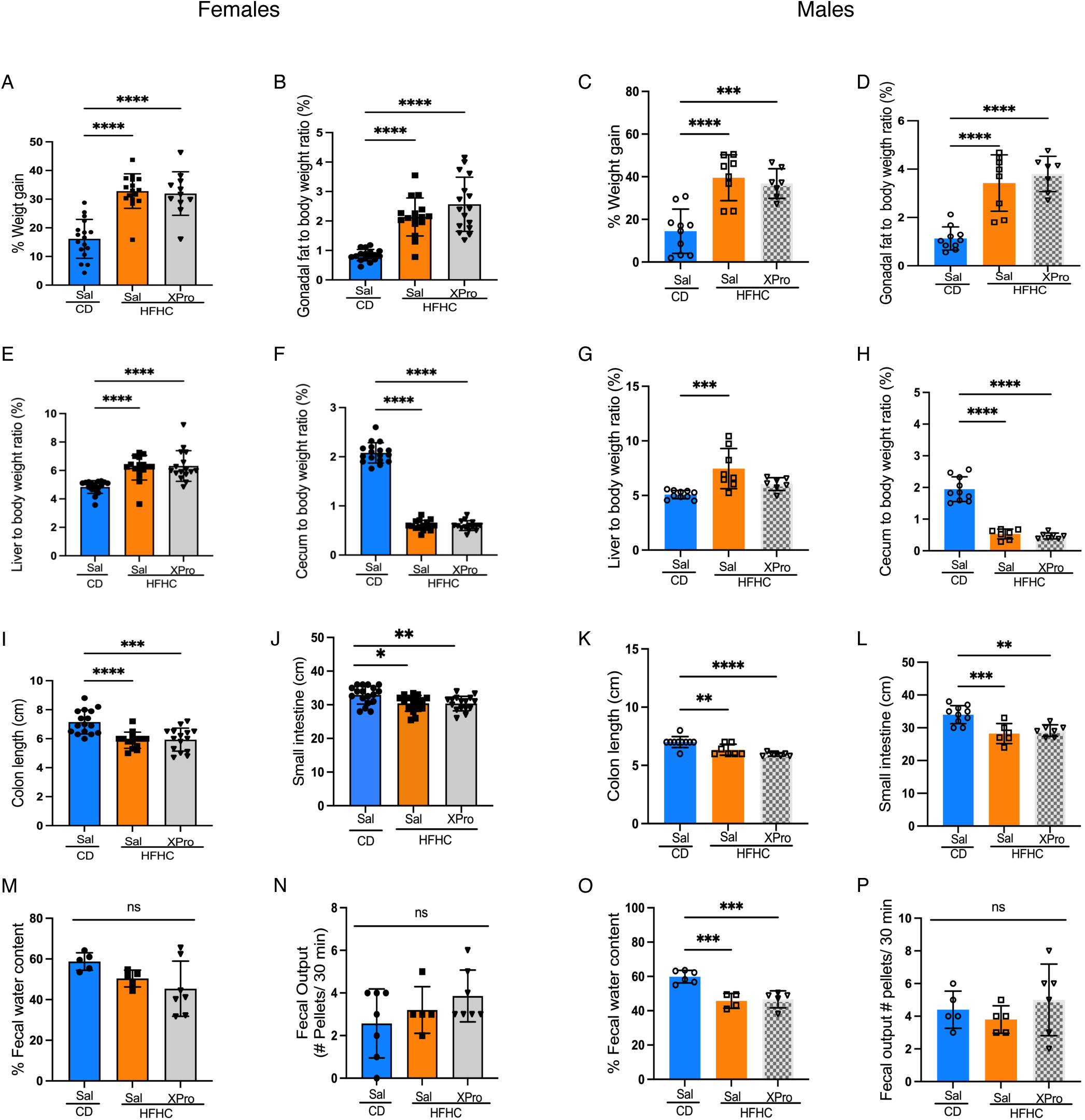
Inhibition of soluble TNF signaling does not rescue HFHC-induced morphometric alterations in 5xFAD mice. (**A-H**) Weight gain, gonadal adipose tissue and liver weights, and cecum size of female and male 5xFAD after 8wks of HFHC treatment. (**I-L**) Diet-induced obesity impact on colon and small intestine lengths on females and male 5xFAD groups. (**M-P**) Fecal water content and fecal output from CD and HFHC groups. Points represent individuals and bars the mean and SEM. Data assessed by one-way ANOVA and Tukey’s post-hoc test. \**p* ≤*0.05*, \*\**p* ≤ 0.01, \*\*\**p* ≤ 0.001, \*\*\*\**p* ≤ 0.0001.

### Soluble TNF inhibition rescues HFHC diet-induced insulinemia and systemic inflammation in male 5xFAD mice

Mounting evidence suggests that solTNF plays a major role in the release of inflammatory cytokines and chemokines that lead to insulin dysfunction and increased risk for AD. While the primary morphometric parameters of HFHC-diet were unchanged following solTNF inhibition, we next tested a contribution of solTNF in metabolic inflammation caused by HFHC diet in 5xFAD mice. Interestingly, XPro1595 had little effect on the circulating metabolic inflammatory mediators tested in female mice (Fig. 4). However, XPro1595 reverted the HFHC-induced impacts on insulin, glucagon, IL-6 and CCL2 in male mice (Fig. 4C, D and G, H). In both sexes, XPro1595 reduced circulating LBP as one measure of intestinal barrier integrity, although only significantly so in female mice (Fig. 4N, P). Altogether, these results suggest that the solTNF signaling contributes to metabolic inflammation induced by a HFHC diet in 5xFAD mice, but in a sex-specific fashion.

**Figure 4.**
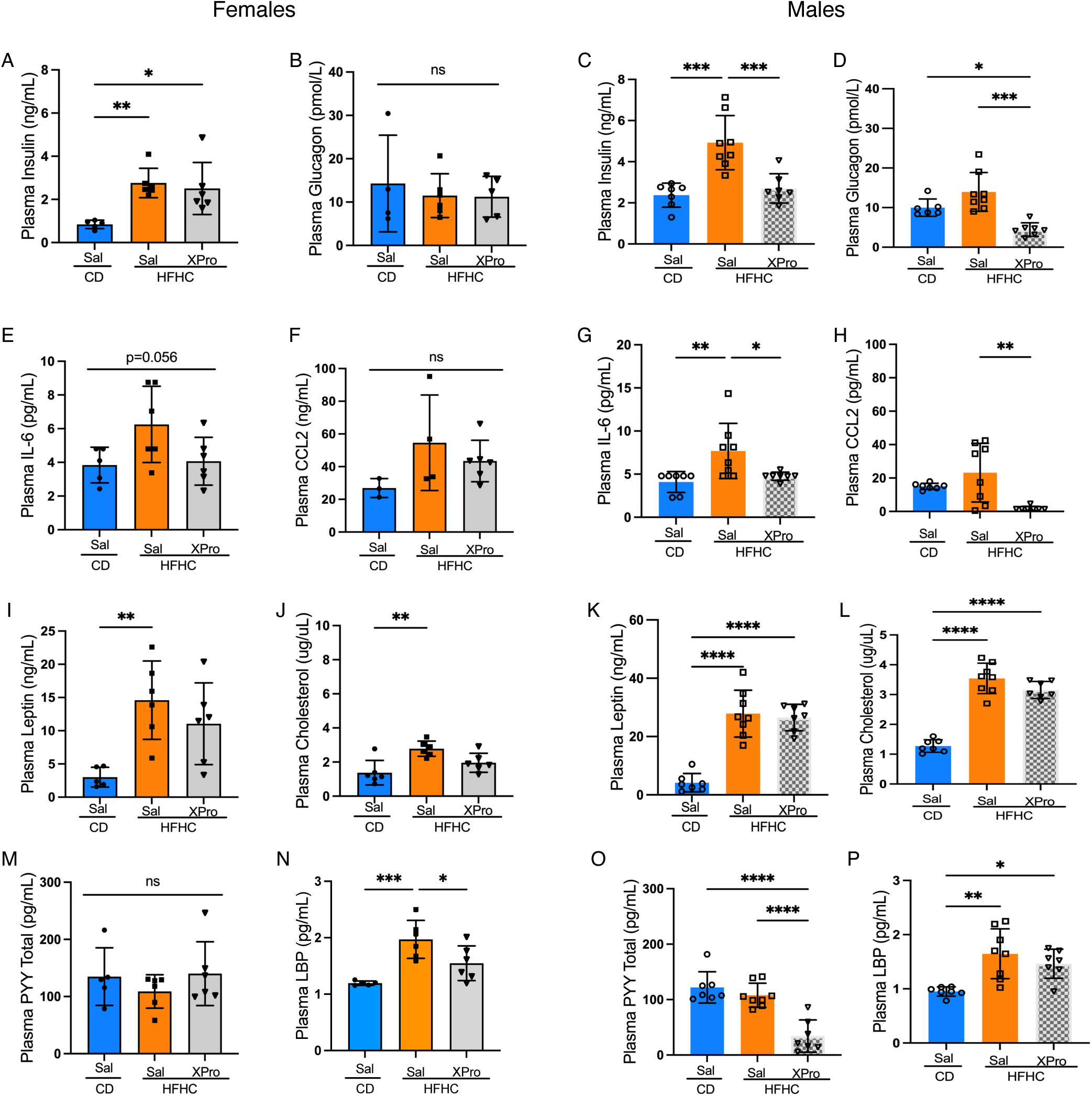
Soluble TNF inhibition rescues HFHC diet-induced insulin impairment and systemic inflammation in male 5xFAD mice. (**A-P**) Circulating concentrations of insulin, glucagon, leptin, cholesterol, IL-6 and CCL2, PYY or LBP in the plasma of male and female 5xFAD mice fed a CD or a HFHC diet for 8 wks and treated with XPro or saline (Sal) for 4wks and determined by multiplexed ELISA. Points represent individuals and bars the mean and SEM. Data assessed by one way ANOVA and Tukey’s post-hoc test. \**p* ≤*0.05*, \*\**p* ≤ 0.01, \*\*\**p* ≤ 0.001, \*\*\*\**p* ≤ 0.0001. ns=non-significant

We therefore further investigated the contribution of solTNF on HFHC-induced inflammation and amyloid parameters, specifically in 5XFAD male mice. We found that XPro1595 reverted diet-induced increases in IL12p70 and IL-10 in colonic tissue, but had no effect on circulating cytokine profiles (Fig. 5A, B). In the CNS, HFHC-fed male 5xFAD mice displayed a decrease in IL-10 in both the hippocampus and cortex, with CXCL2 also decreased in the cortex and reversed by solTNF inhibition (Fig. 5C-F). Male animals also displayed an increase in soluble Aβ42 and Aβ40 in the hippocampus after 8 weeks of HFHC intake (Fig. 6A-D). However, HFHC diet did not induce notable alterations to the Aβ40/42 ratio or Aβ38, nor did solTNF inhibition revert Aβ concentrations (Fig. 6A-D). XPro1595 did result in substantially decreased both soluble Aβ42 and 40 in plasma, and increased the plasma Aβ40/42 ratio (Fig. 6E, F). These findings suggest that the systemic and central metabolic and immune interactions in response to a HFHC diet promote a neurotoxic hippocampal in male 5xFAD mice.

**Figure 5.**
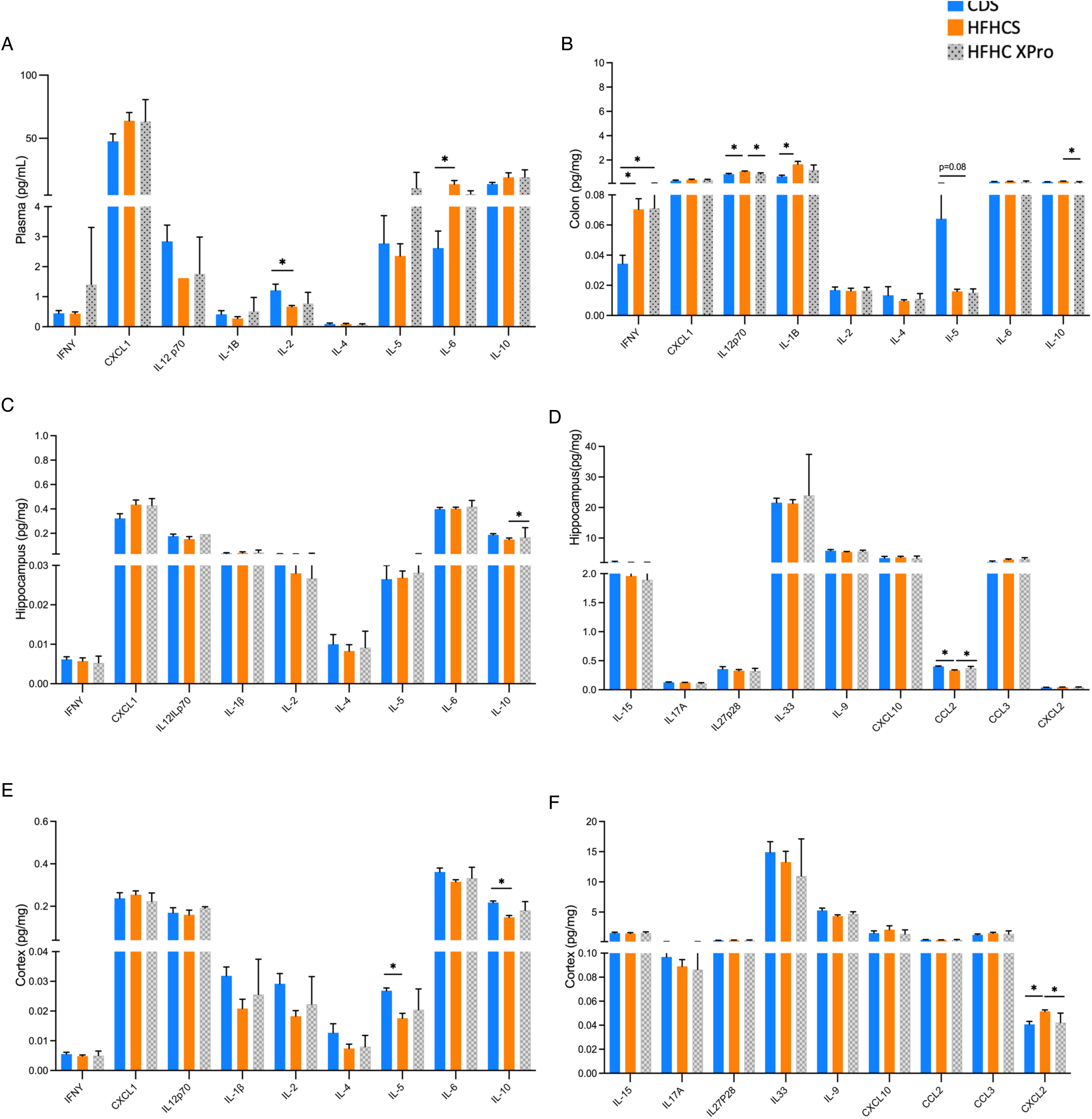
Soluble TNF inhibition mitigates HFHC-induced immune alterations. Multiplexed ELISA analysis of cytokine and chemokine concentrations in the (A) plasma, (B) colon, (C, D) hippocampus and (E, F) cortex of 5xFAD male mice after 8wks of a CD (CDS) or HFHC consumption and XPro treatment (HFHC XPro) or vehicle (HFHCS). Bars represent the mean and SEM. Data assessed by one-way ANOVA with Tukey’s post-hoc test. \**p* ≤*0.05*; n=5-10.

**Figure 6.**
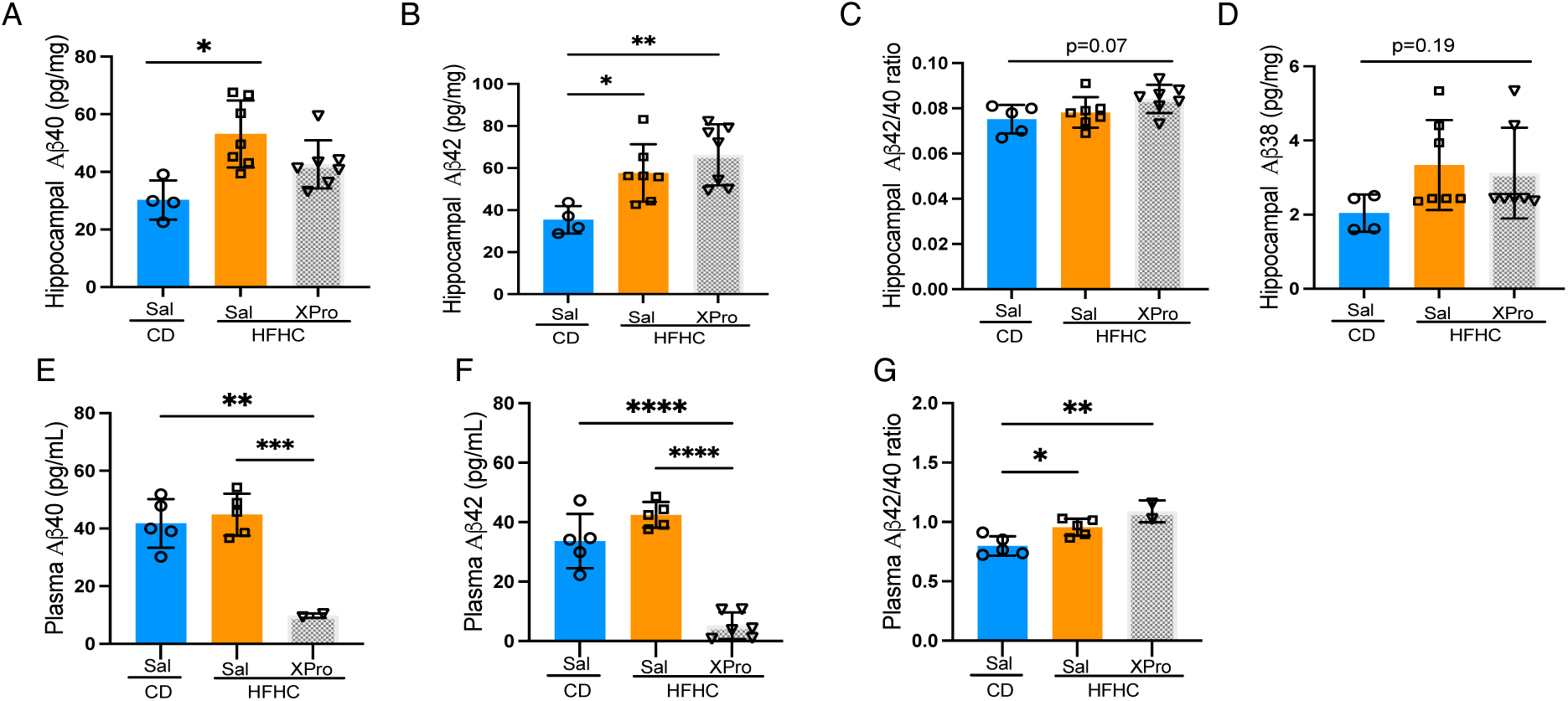
Amyloid beta increases following HFHC-diet in male 5xFAD mice. Multiplexed ELISA analysis from the (**A-D**) hippocampus or (**E-G**) plasma of male 5xFAD mice fed CD or HFHC diets for 8wks and subsequent treatment with XPro or vehicle (Sal). Points represent individuals and bars the mean and SEM. Data assessed by one-way ANOVA with Tukey’s post-hoc test. \**p* ≤*0.05*, \*\**p* ≤ 0.01, \*\*\**p* ≤ 0.001, \*\*\*\**p* ≤ 0.0001.

### solTNF signaling contributes to HFHC-induced transcriptional state in the CNS

Given the effects on metabolic and inflammatory signaling in both circulation and the CNS, we set out explore broader impacts of both HFHC diet and solTNF signaling on the CNS transcriptional landscape. Targeted transcriptomics via Nanostring assay revealed differential gene expression patterns in the hippocampus and cortex in male and female mice, in response to both diet and XPro1595 treatment (Fig. 7 and Supplemental Tables 1-8). Qualitatively, female 5xFAD mice displayed an increased frequency of differentially expressed genes, in both diet and solTNF interventions, compared to identically treated male animals. Further, we note little to no overlap between differentially expressed genes between either brain region or by sex, highlighting regional and sex-specificity to the effects of HFHC diet and XPro1595 inhibition. Generally, we observe that HFHC promotes a more inflammatory transcriptional state in female 5xFAD mice, but that genes associated with amyloid deposition appear upregulated in HFHC-fed male 5xFAD animals. XPro1595 treatment and solTNF inhibition does not fully downregulate the same profile of genes that HFHC-diet induces; however we note that it increases hippocampal choline acetyltransferase (ChAT) expression in female mice, perhaps indicative of neuroprotective pathways. Thus, in line with the metabolic and inflammatory profiling, HFHC diet influences AD-relevent neuroinflammatory pathways in the CNS, partially mediated by solTNF signaling.

**Figure 7.**
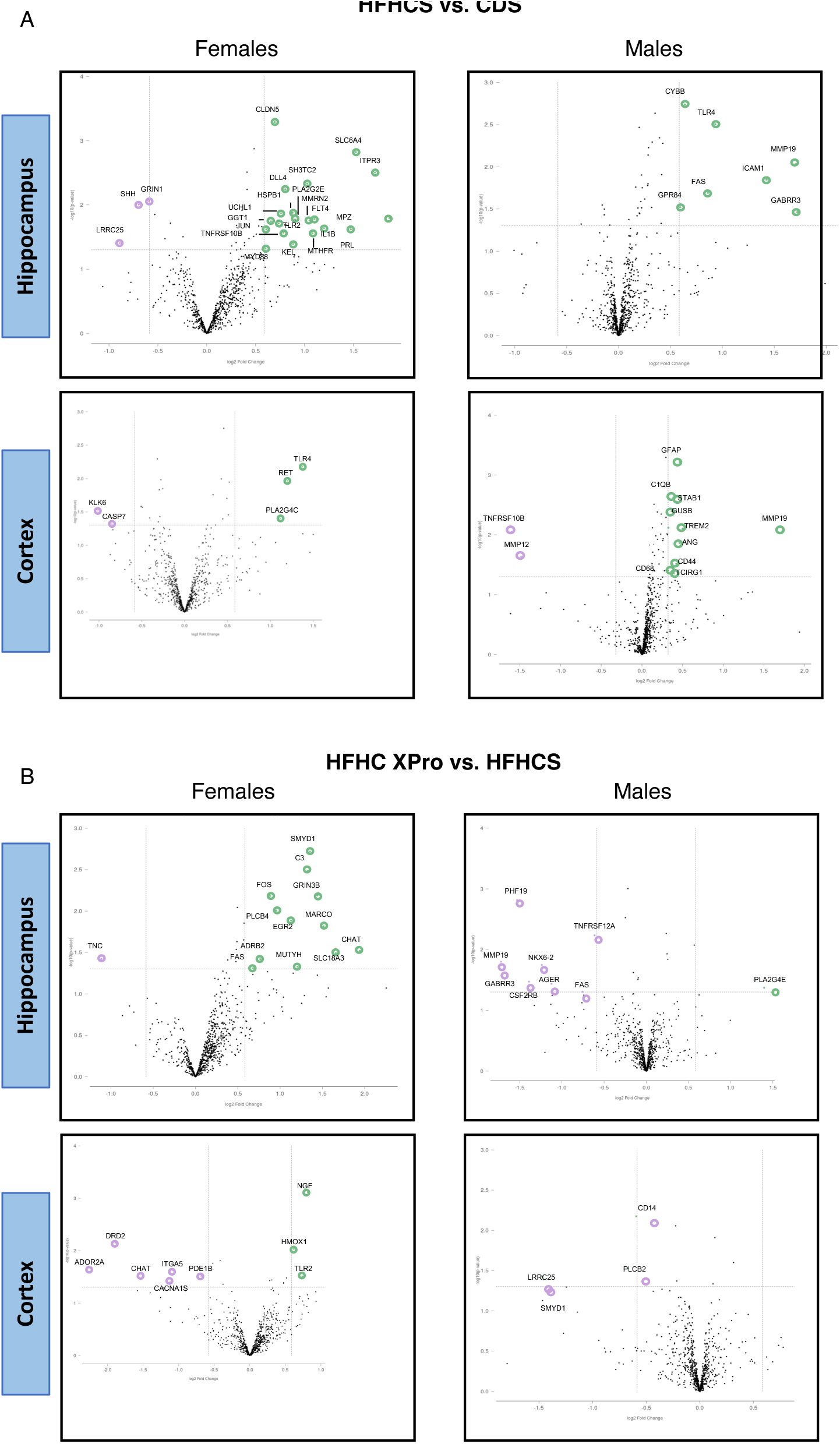
HFHC diet and solTNF signaling differentially impact CNS transcriptional status in a sex-specific fashion. Volcano plots depicting differences in expression across indicated comparison groups, as determined by Nanostring targeted transcriptional profiling. (**A**) Diet-dependent gene expression or (**B**) solTNF-dependent gene expression in hippocampal and cortical tissue from male and female 5xFAD mice after 8wks of a CD (CDS) or HFHC consumption treated with vehicle (HFHCS) or XPro (HFHC XPro). Highlighted genes represent those with a fold-change >[1.5] and unadjusted *p* value of 0.05. N=5 per group.

## Discussion

Metabolic disorders associated with obesity, such as type 2 hypercholesterolemia and diabetes, are marked by metabolic inflammation, which greatly increases risk for dementia (35, 43, 49). Metabolic inflammation is a mild, yet chronic and systemic inflammation that involves intricate interactions between adipose tissue, liver, the microbiome, and intestine in response to high-energy diet consumption (50, 51). Immune activation in obesity and sustained inflammatory activity impact the CNS through the persistent release of cytokines and chemokines (52). Recent research has shown that systemic inflammation contributes to AD pathologies through several mechanisms, including the increase of amyloid deposition and the promotion of a neurotoxic environment (35, 53). Here, we show that the gene expression and immune responses to an HFHC diet in 5xFAD mouse model of AD are influenced by compartment, sex and solTNF signaling.

Immune dysregulation plays a crucial role in both obesity and metabolic syndrome, and AD development and progression. For instance, the glycogen synthase kinase 3 beta (GSK3β) is a metabolic regulatory molecule often associated with insulin dysfunction that is hyperactivated in the CNS of those with AD and animal models of dementia (47, 54, 55). GSK-3β can activate NF-κB which results in further cytokine and chemokine responses (56). In fact, here we found that a HFHC diet increases hippocampal GSK-3β in female 5xFAD mice demonstrating the activation of this pathway by diet in this AD mouse model. A recent study that investigated shared genes between AD and T2D suggested that IL-1β, TNF and IL-10 are common signaling molecules that connect both diseases (57). Correspondingly, we observe similar alterations with HFHC diet inducing decreased IL-10 and increased IL-1 β in the CNS without further evidence of inflammasome activation or impacts on Aβ abundance. These data suggest both specificity to the inflammatory impacts of HFHC diet and highlights the potential effects of these on pathways activated prior to overt neurodegeneration and amyloid deposition.

Our team previously showed that a HFHC diet can modulate CNS-associated CD8+ T cell populations, without a significant impact on amyloidogenesis, in this same 5xFAD model of AD (42). Here, our findings indicate that the HFHC-induced CNS pro-inflammatory state corresponds with systemic metabolic and immune dysregulation, largely in a sex-specific fashion. In agreement with that prior work, despite the significant CNS inflammatory environmental alterations present following HFHC diet in female mice, no detectable changes in Aβ40 and Aβ42 in this early age was observed with this diet regimen. Conversely, while male mice appeared less robustly inflammatory following HFHC diet, Aβ abundances increased, highlighting sex-specific impacts of metabolic dysregulations on these two arms of AD-related pathologies. We also note that solTNF signaling, modulated by XPro1595, did not appear to contribute robustly to HFHC-induced metabolic parameters in female 5xFAD mice, but did have a significant effect on male animals. Prior work from our team demonstrated a beneficial role of solTNF inhibition in reverting insulinemia and systemic inflammation in an obesogenic environment in wild-type male mice (58). Our new data herein indicate a sex-dependent impact of solTNF inhibition in reverting increases in circulating insulin and inflammatory markers (CCL2 and IL-6) in this model of AD pathologies. Others have also observed sex disparities in in immune responses in the 5xFAD mice. A recent study demonstrated that female 3xTG-AD mice are more susceptible to metabolic alterations in response to a HF diet (59, 60). Our findings in this study suggest that the solTNF signaling may contribute centrally and systemically in regulating metabolic inflammation both in wildtype genetic environments and genetically-predisposed models of AD in a sex-specific manner.

Interestingly, we observed distinct cytokine and metabolic profiles induced by HFHC diet across compartments. However, some shared features emerged. For instance, HFHC induced a decrease in IL-5 and an increase in CXCL2 and IL-1β in the colon, alongside similar alterations in CNS tissues. Impacts to the immune environment co-occurred with a compositional shift of the gut microbiome, which may facilitate release of pro-inflammatory mediators and decreases in occluding production which may allow those microbial molecules to pass more readily into circulation. In line with this, we note that HFHC induces an increase in circulating LBP, suggesting a response to microbial-derived molecules. Our team and others have observed similar effects in prior work, both in the context of insulin dysregulation (57, 61, 62).

While there is a general understanding about the connection between immune alterations and AD, several questions remain regarding the interaction between central and peripheral immune system responses in the context of modifiable factors, such as an unhealthy diet. Hormones that regulate energy metabolism appear as a possible link to connect these systemic and central immune actions in dementia (63). In this study, we provide a foundation to begin parsing out the intricate relationship between sex, obesity, and AD-related pathologies in an environment characterized by solTNF. Furthermore, we have shown the beneficial impact of inhibiting solTNF on the metabolic and immune interplay in metabolic inflammation and AD, and uncovered sex-specific outcomes to this signaling cascade. These data emphasize the necessity for comprehensive research to explore potential drug-intervention interactions involving metabolic and sex-dependent pathways for future avenues of therapeutics.

## Ethics declarations

All animal husbandry and experiments were approved by the Institutional Animal Care and Use Committee of Emory University, #PROTO201900056.

## Author Contributions

Maria Elizabeth de Sousa Rodrigues: Conceptualization, methodology, investigation, analysis, Data curation, original draft preparation, writing, reviewing and editing, funding acquisition MacKenzie Bolen: Investigation, analysis, data curation, writing, reviewing and editing

Lisa Blackmer-Raynolds: Investigation, writing, reviewing and editing Noah Schwartz: Investigation, reviewing and editing

Jianjun Chang: Investigation, reviewing and editing

Malù Gàmez Tansey: Funding acquisition, conceptualization, reviewing and editing

Timothy Robert Sampson: Funding acquisition, original draft preparation, writing, reviewing and editing.

## Acknowledgments

We thank members of the Sampson lab for productive discussions and Isabel Fraccaroli for administrative and technical assistance. We acknowledge support from the Emory Integrated Genomics Core (EIGC), the Emory Multiplexed Immunoassay Core (EMIC), and the Division of Animal Resources (DAR) at Emory University - Integrated Core Facilities of the Emory University School of Medicine and supported by the Georgia Clinical and Translational Science Alliance of the NIH (UL1TR002378). Dr. David Szymkowski at Xencor, Inc. for providing XPro1595. Funding for this work was derived from the National Institutes of Aging/National Institutes of Health grants RF1AG051514 (MGT), RF1AG057247 (MGT), F31AG076332 (LBR) and an Alzheimer’s Association Research Fellowship to Promote Diversity program (AARFD-19-619290; MESR).

## Supplemental Figure Legends

**Supplemental Figure S1.**
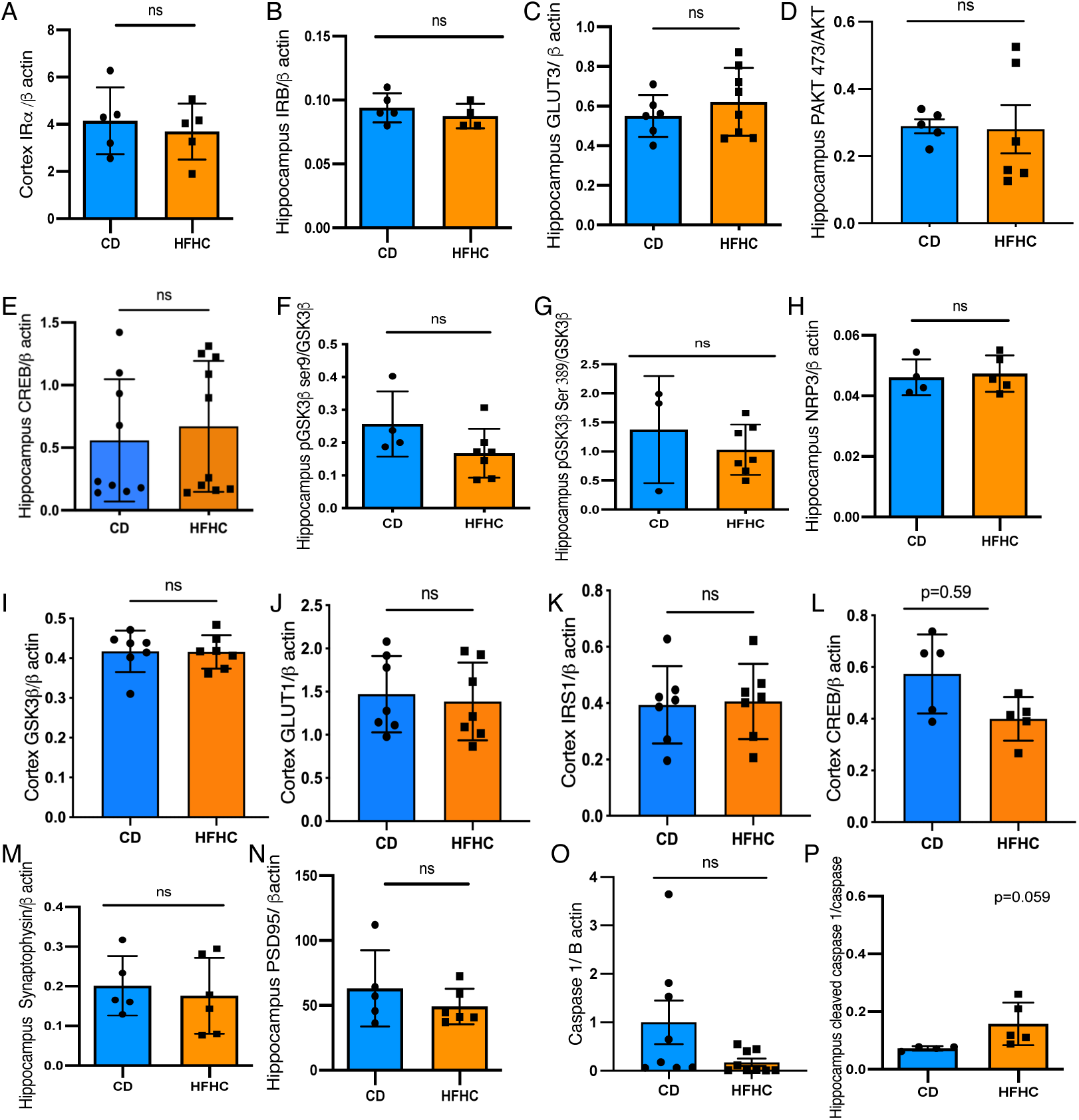
High-fat high-carbohydrate impact on metabolic pathways in the hippocampus and cortex of 5xFAD female mice. Densitometric quantification of western blot analysis of hippocampal and cortical tissue from female 5xFAD mice fed 8wks of a CD or HFHC, for the indicated metabolic signaling proteins. Points represent individuals and bars the mean and SEM. Data assessed by unpaired, two-tailed T-test. \**p* ≤*0.05*, \*\**p* ≤ 0.01, \*\*\**p* ≤ 0.001, \*\*\*\**p* ≤ 0.0001.

**Supplemental Figure S2.**
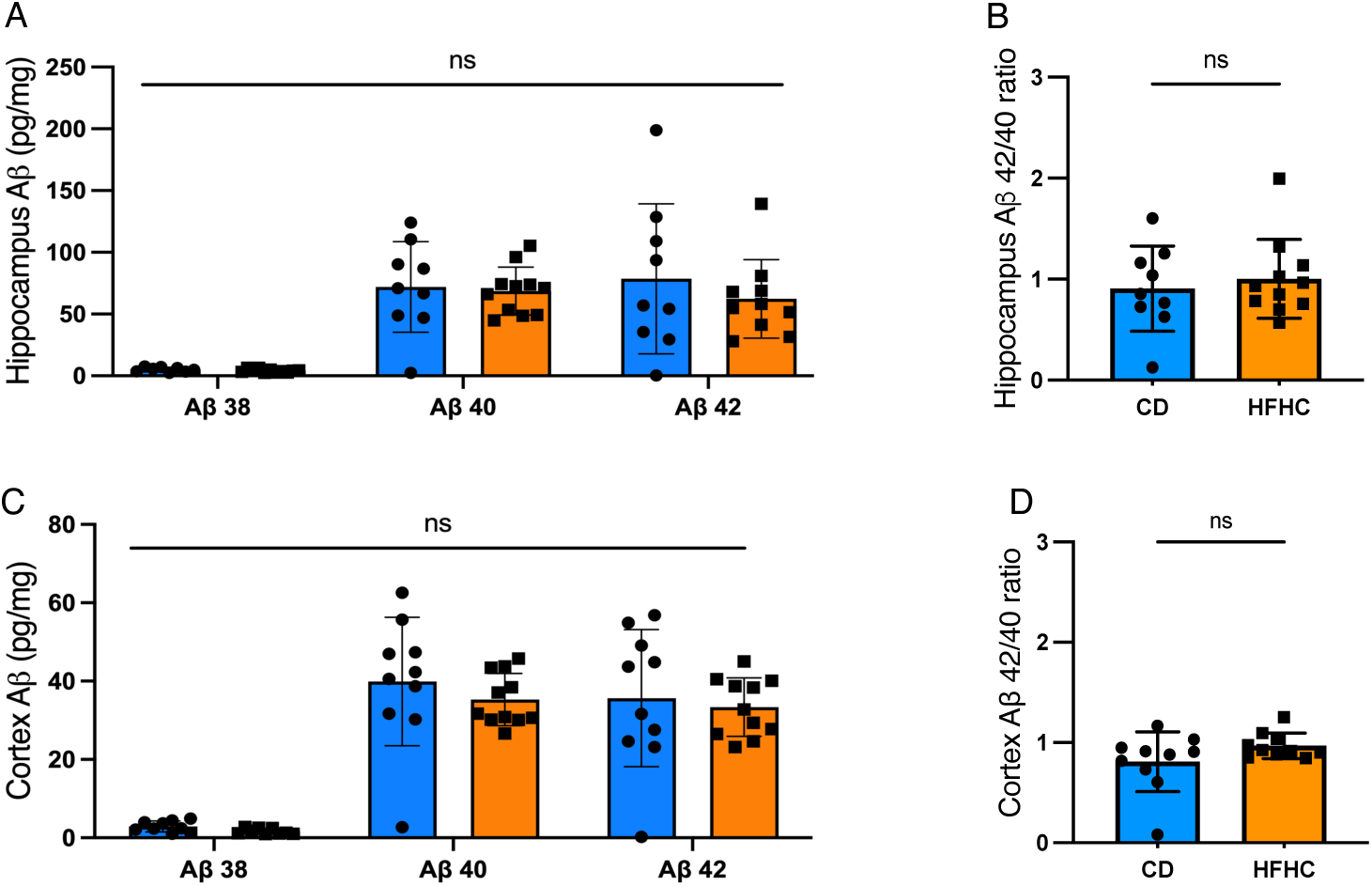
Amyloid beta abundance is not altered by HFHC in female 5xFAD mice. (**A-D**) Aβ 38, 40 and 42 in soluble homogenates from (**A, B**) hippocampal and (**C, D**) cortical tissue quantified by multiplex immunoassays from 5xFAD mice after 8wks of CD or HFHC consumption. Points represent individuals and bars the mean and SEM. Data assessed by unpaired, two-tailed T-test. ns=non-significant.

**Supplemental Figure S3.**
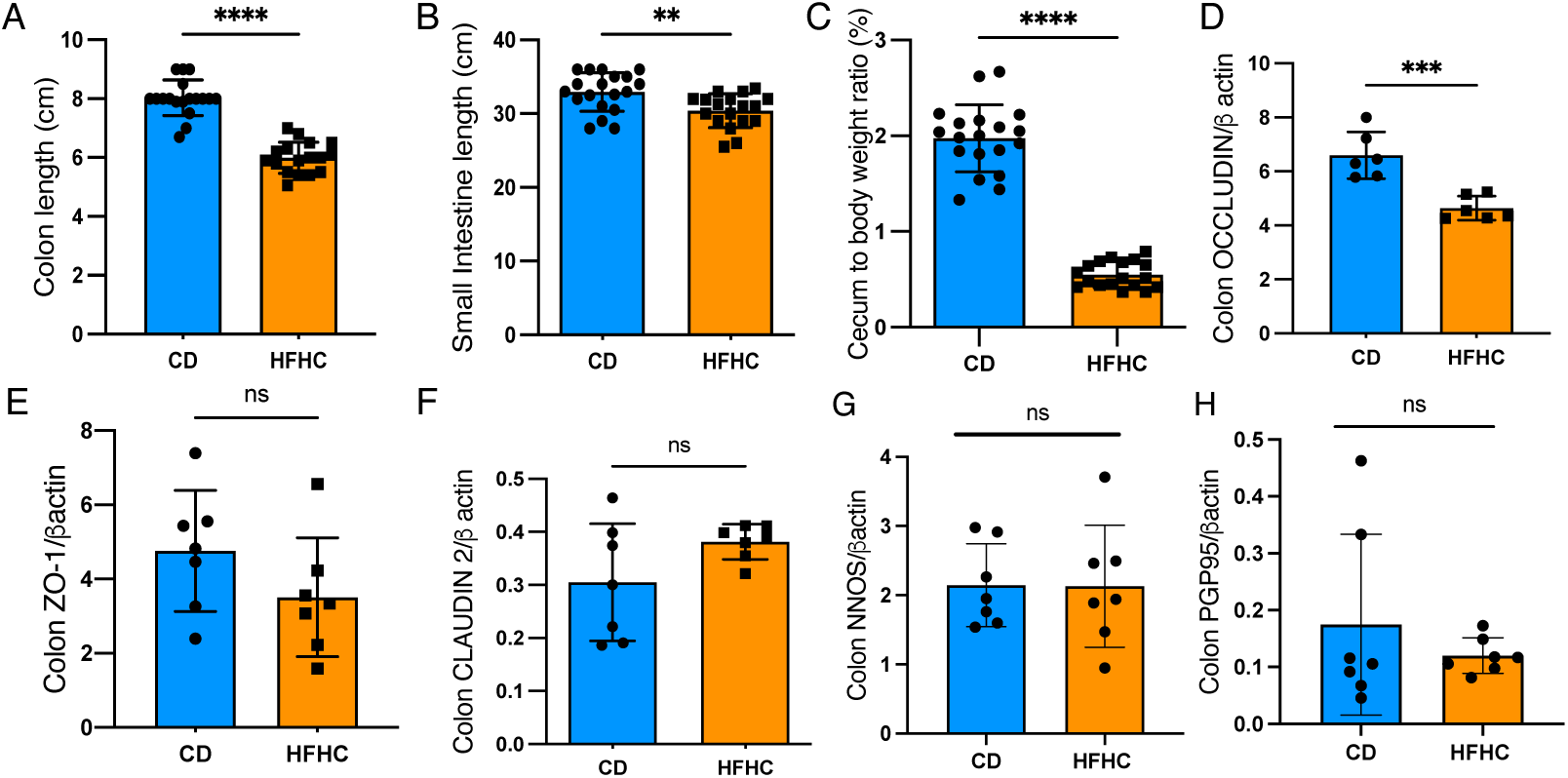
High-fat high-carbohydrate intake is associated with intestinal dysregulation. (**A**) Colon length, (**B**) small intestine length, and (**C**) cecum weight of female 5xFAD mice fed 8wks of a CD or HFHC. (**D-H**) Western blot analysis of proximal colon for occludin, ZO-1, claudin 2, nNOS and PGP9.5. Points represent individuals and bars the mean and standard error. Data assessed by unpaired, two-tailed T-test. \*\**p* ≤ 0.01, \*\*\**p* ≤ 0.001, \*\*\*\**p* ≤ 0.0001.

**Supplemental Figure S4.**
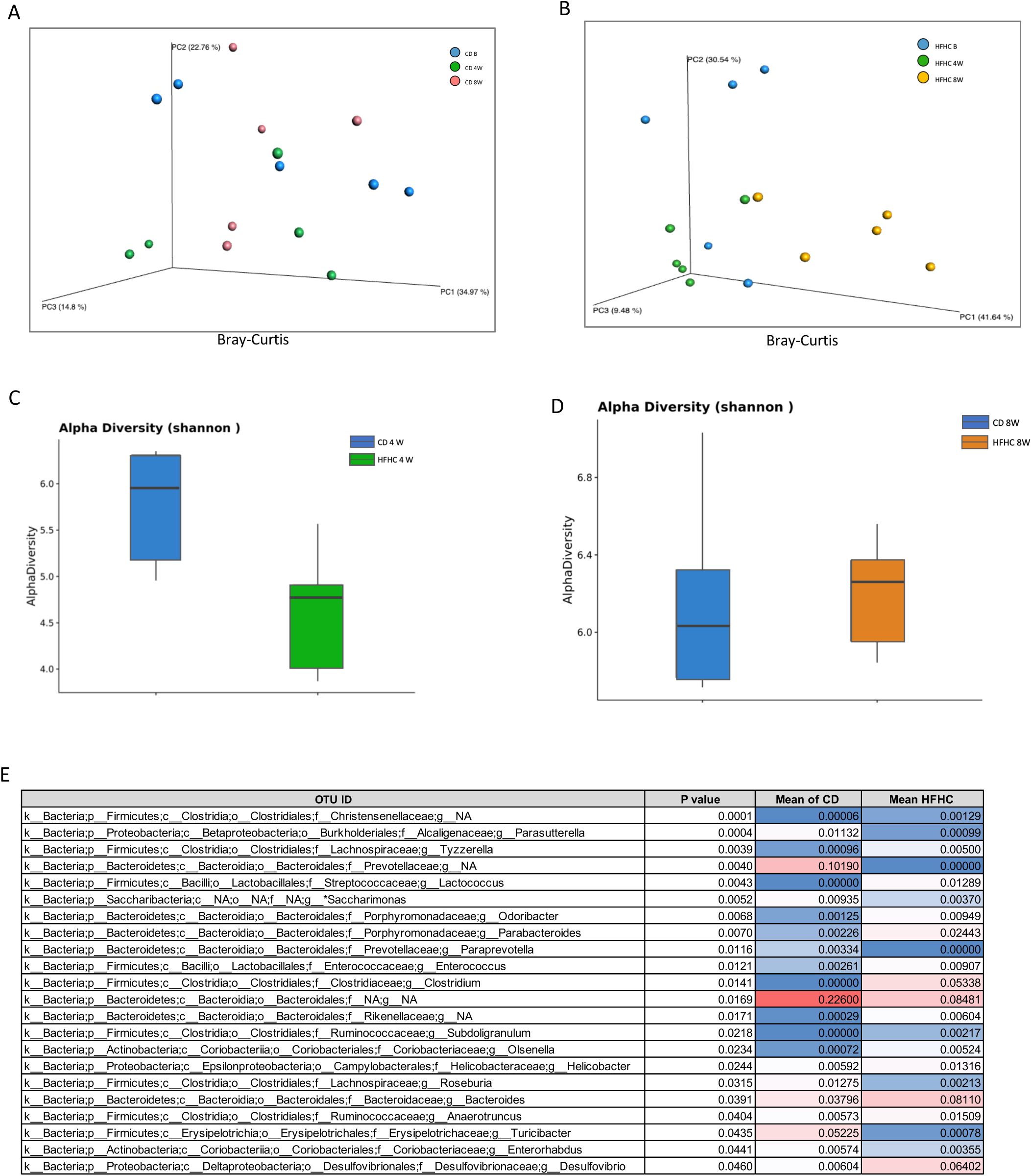
Diet-induced obesity impact on intestinal microbiome in 5xFAD mice. (**A**) PCA plots based on Bray-Curtis distance of 16S microbiome profiles from mice receiving the CD or (**B**) HFHC over 8wks. (**C, D**) Shannon alpha diversity indices comparing intestinal microbial community between 5xFAD fecal samples from mice fed a CD or HFHC diet at the (**C**) 4 wks or (**D**) 8 wks time points post-diet intervention. (**E**) Genera-level alterations between diets at 8wks, determined by two-tailed T test. Points represent individuals (**A, B**), and boxes the range (**C, D**). N=5 from individual cages.

