## Supplemental Tables for "Diet-induced metabolic and immune impairments are sex-specifically modulated by soluble TNF signaling in the 5xFAD mouse model of Alzheimer’s disease"

**Supplemental Table 1.** Hippocampal tissue from female mice comparing HFHC-diet, Saline-treated to Control diet, Saline treated.

| Gene | *p* value | Fold change | Adj *p* value |
|---|---|---|---|
| Grin1 | 0.0087 | -1.5 | 0.65 |
| Lrrc25 | 0.039 | -1.9 | 0.66 |
| Shh | 0.0098 | -1.6 | 0.65 |
| Cldn5 | 0.00051 | 1.7 | 0.38 |
| Dll4 | 0.006 | 1.7 | 0.55 |
| Flt4 | 0.017 | 2.1 | 0.65 |
| Ggt1 | 0.018 | 1.58 | 0.65 |
| Hspb1 | 0.013 | 1.8 | 0.65 |
| Il1b | 0.023 | 2.3 | 0.65 |
| Itpr3 | 0.0031 | 3.3 | 0.47 |
| Jun | 0.024 | 1.5 | 0.65 |
| Kel | 0.041 | 1.85 | 0.66 |
| Mmrn2 | 0.017 | 2.07 | 0.65 |
| Mpz | 0.024 | 2.8 | 0.65 |
| Mthfr | 0.028 | 2.1 | 0.66 |
| Myd88 | 0.048 | 1.5 | 0.66 |
| Pla2g2e | 0.017 | 1.9 | 0.65 |
| Prl | 0.016 | 1.87 | 0.65 |
| Sh3tc2 | 0.0046 | 2 | 0.54 |
| Slc6a4 | 0.0015 | 2.9 | 0.38 |
| Tlr2 | 0.019 | 1.7 | 0.65 |
| Tnfrsf10b | 0.027 | 1.7 | 0.66 |
| Uchl1 | 0.013 | 1.7 | 0.65 |

**Supplemental Table 2.** Hippocampal tissue from female mice comparing HFHC-diet, XPro-treated to HFHC-diet, Saline treated.

| Gene | *p* value | Fold change | Adj *p* value |
|---|---|---|---|
| Tnc | 0.038 | -2.16 | 0.97 |
| ADRB2 | 0.038 | 1.7 | 0.97 |
| C3 | 0.003 | 2.51 | 0.97 |
| Chat | 0.03 | 3.84 | 0.97 |
| Egr2 | 0.013 | 2.19 | 0.97 |
| Fas | 0.049 | 1.6 | 0.97 |
| Fos | 0.0066 | 1.84 | 0.97 |
| Grin3b | 0.0065 | 2.72 | 0.97 |
| Marco | 0.015 | 2.85 | 0.97 |
| Mutyh | 0.047 | 2.31 | 0.97 |
| Plcb4 | 0.0099 | 1.94 | 0.97 |
| Slc18a3 | 0.032 | 3.16 | 0.97 |
| Smyd1 | 0.0019 | 2.55 | 0.97 |

**Supplemental Table 3.** Hippocampal tissue from male mice comparing HFHC-diet, Saline-treated to Control diet, Saline treated.

| Gene | *p* value | Fold change | Adj *p* value |
|---|---|---|---|
| Cybb | 0.0018 | 1.56 | 0.56 |
| Fas | 0.02 | 1.82 | 0.86 |
| Gabrr3 | 0.034 | 3.3 | 0.93 |
| Gpr84 | 0.03 | 1.51 | 0.93 |
| Icam1 | 0.014 | 2.68 | 0.79 |
| Mmp19 | 0.0093 | 3.26 | 0.71 |
| Tlr4 | 0.0031 | 1.91 | 0.56 |

**Supplemental Table 4.** Hippocampal tissue from male mice comparing HFHC-diet, XPro-treated to HFHC-diet, Saline treated.

| Gene | *p* value | Fold change | Adj *p* value |
|---|---|---|---|
| Ager | 0.037 | -2.18 | 0.97 |
| Csf2rb | 0.034 | -2.62 | 0.97 |
| Fas | 0.05 | -1.69 | 0.97 |
| Gabrr3 | 0.021 | -3.21 | 0.97 |
| Mmp19 | 0.016 | -3.3 | 0.97 |
| Nkx6-2 | 0.018 | -2.36 | 0.97 |
| Phf19 | 0.0016 | -2.89 | 0.58 |
| Tnfrsf12a | 0.0058 | -1.53 | 0.88 |
| Pla2g4e | 0.042 | 2.63 | 0.97 |

**Supplemental Table 5**. Frontal cortex tissue from female mice comparing HFHC-diet, Saline-treated to Control diet, Saline treated.

| Gene | *p* value | Fold change | Adj *p* value |
|---|---|---|---|
| Casp7 | 0.048 | -1.8 | 0.98 |
| Klk6 | 0.031 | -2 | 0.98 |
| Pla2g4c | 0.04 | 2.2 | 0.98 |
| Ret | 0.011 | 2.3 | 0.98 |
| Tlr4 | 0.0066 | 2.6 | 0.98 |

**Supplemental Table 6.** Frontal cortex tissue from female mice comparing HFHC-diet, XPro-treated to HFHC-diet, Saline treated.

| Gene | *p* value | Fold change | Adj *p* value |
|---|---|---|---|
| Adora2a | 0.022 | -4.69 | 1 |
| Cacna1s | 0.04 | -2.18 | 1 |
| Chat | 0.032 | -2.85 | 1 |
| Drd2 | 0.0071 | -3.71 | 1 |
| Itga5 | 0.024 | -2.12 | 1 |
| Pde1b | 0.03 | -1.61 | 1 |
| Hmox1 | 0.0093 | 1.54 | 1 |
| Ngf | 0.00077 | 1.76 | 0.59 |
| Tlr2 | 0.031 | 1.67 | 1 |

**Supplemental Table 7**. Frontal cortex tissue from male mice comparing HFHC-diet, Saline-treated to Control diet, Saline treated.

| Gene | *p* value | Fold change | Adj *p* value |
|---|---|---|---|
| Mmp12 | 0.023 | -2.79 | 0.56 |
| Tnfrsf10b | 0.009 | -3.04 | 0.39 |
| Ang | 0.014 | 1.38 | 0.43 |
| C1qb | 0.0024 | 1.31 | 0.38 |
| C1qc | 0.0076 | 1.25 | 0.39 |
| Cd44 | 0.03 | 1.34 | 0.67 |
| Cd68 | 0.042 | 1.29 | 0.78 |
| Gfap | 0.00061 | 1.38 | 0.23 |
| Gusb | 0.004 | 1.29 | 0.38 |
| Mmp19 | 0.0084 | 3.32 | 0.39 |
| Stab1 | 0.0028 | 1.35 | 0.38 |
| Tcirg1 | 0.044 | 1.31 | 0.79 |
| Trem2 | 0.0079 | 1.42 | 0.39 |

**Supplemental Table 8.** Frontal cortex tissue from female mice comparing HFHC-diet, XPro-treated to HFHC-diet, Saline treated.

| Gene | *p* value | Fold change | Adj *p* value |
|---|---|---|---|
| Cd14 | 0.0067 | -1.51 | 1 |
| Lrrc25 | 0.038 | -2.76 | 1 |
| Plcb2 | 0.03 | -1.59 | 1 |
| Smyd1 | 0.041 | -2.75 | 1 |
